# Deep learning based brain age prediction uncovers associated sequence variants

**DOI:** 10.1101/595801

**Authors:** B.A. Jonsson, G. Bjornsdottir, T.E. Thorgeirsson, L.M. Ellingsen, G. Bragi Walters, D.F. Gudbjartsson, H. Stefansson, K. Stefansson, M.O. Ulfarsson

**Author notes:** Corresponding author Email addresses (K. Stefansson), (M.O. Ulfarsson).

## Abstract

Machine learning algorithms trained to recognize age-related structural changes in magnetic resonance images (MRIs) of healthy individuals can be used to predict biological brain age in independent samples. The difference between predicted and chronological age, predicted age difference (PAD), is a phenotype holding promise for the study of normal brain ageing and brain diseases, and genetic discovery *via* genome-wide association studies (GWASs). Here, we present a new deep learning approach to predict brain age from a T1-weighted MRI. The method was trained on a dataset of healthy Icelanders (*N* = 1264) and tested on two datasets, the IXI (*N* = 544) and UK Biobank (*N* = 12395) datasets, utilizing transfer learning to improve accuracy on new sites. A GWAS of PAD in the UK Biobank data (discovery set: N=12395, replication set: N=4453) yielded two sequence variants, rs1452628-T (*β*=-0.08, *P* = 1.15 · 10^−9^) and rs2435204-G (*β*=0.102, *P* = 9.73 · 10^−12^). The former is near *KCNK2* and correlates with reduced sulcal width, whereas the latter correlates with reduced white matter surface area and tags a well-known inversion at 17q21.31 (H2). The genetic association analysis was also confined to variants known to associate with brain structure, yielding three additional sequence variants associating with PAD.

## 1. Introduction

Ageing has a significant structural impact on the brain that correlates with decreased mental and physical fitness [12] and increased risk of neurodegenerative diseases such as Alzheimer’s disease [1] and Parkinson’s disease [59]. Recent publications, have demonstrated that MRIs can be used to predict chronological age with reasonably good accuracy [20, 49, 12]. Such predictions provide an estimate of biological brain age in independent samples. The traditional way to perform brain age prediction is to extract features from brain MRIs followed by classification or regression analysis. This includes extracting principal components [20], cortical thickness and surface curvature [77], volume of gray matter (GM), white matter (WM), and cerebrospinal fluid [38], and constructing a similarity matrix [10]. The drawback of using feature extraction methods is loss of information since the features are likely not designed explicitly for extracting information relevant to brain age. Recently, deep learning (DL) methods have garnered much interest [46]. These methods learn features that are important without apriori bias or hypothesis. Convolutional neural networks (CNNs) [45] are deep learning techniques that are especially powerful for image processing and computer vision. Previously, they have been applied to brain age prediction [32, 11]. Notably, Cole et al. [11] implemented a 3D CNN trained on T1-weighted MRIs to predict brain age and achieved promising results.

PAD (the difference between predicted brain age and chronological age) estimates the deviation from healthy ageing. Studies have shown that positive PAD correlates with measures of reduced mental and physical fitness; including weaker grip strength, poorer lung function, slower walking speed, lower fluid intelligence, higher allostatic load and increased mortality risk [12]. In addition, positive PAD has been shown to associate with cognitive impairments [21, 24, 10, 49], diabetes [22], traumatic brain injuries [10], schizophrenia [41, 64], and chronic pain [43]. On the other hand, a negative PAD associates with higher educational attainment [72], increased physical activity [72] and meditation [50]. Moreover, PAD has been demonstrated to be heritable [11, 35] and to have a polygenic overlap with brain disorders such as schizophrenia, bipolar disorder, multiple sclerosis, and Alzheimer’s disease [35]. Furthermore, the high degree of genetic correlation found among psychiatric and some neurological disorders suggests that current diagnostic boundaries do not necessarily reflect underlying biology [14]. Hence, defining a novel phenotype capturing global age-related changes in brain structure could, *via* variants in the sequence of the genome that associate with these changes, provide novel biological insights.

Here we present a new brain age prediction method (Figure 1A) that uses a 3D CNN trained on MRIs to predict brain age. The input data are a T1-weighted image registered to Montréal Neurological Institute (MNI) space and data derived from the T1-weighted image, i.e., a Jacobian map, and gray and white matter segmented images (Figure 1B). The input data also include information about the subject’s sex and the type of MRI scanner. The output of the network is the predicted brain age.

As mentioned above, Cole et al. [11] trained a 3D CNN to perform brain age prediction. Our network is different in four key ways. **1)** We use a significantly different architecture. While their architecture resembles a standard VGGNet architecture [66] our architecture uses the recent ResNet design [26]. One of the drawbacks of the VGG architecture is that the vanishing gradient problem limits the potential depth of the network. In contrast, the ResNet architecture has no such depth limits. ResNets also have smoother loss surfaces [48], which in turn helps speeding up convergence. **2)** We add inputs to the final CNN layer to factor in information about sex and scanner. **3)** Our technique is the first to use deformation information encoded in Jacobian maps to predict brain age. **4)** As we have mentioned, our method combines predictions from multiple CNNs by either averaging predictions or by training a data blender.

In experiments, we compare our proposed method to a few brain age prediction methods based on feature extraction and machine learning. We also demonstrate that transfer learning is useful for adapting a CNN trained to predict brain age on one site to a new site while retaining predictive accuracy. And we look at how the PAD calculated with our method is affected by random weight initialization and retraining.We then check for associations between PAD and performance on neuropsychological tests. Finally, we perform genetic analysis on PAD using UK Biobank data, resulting in identification of associations with five sequence variants for which we provide detailed phenotypic characterizations.

**Figure 1:**
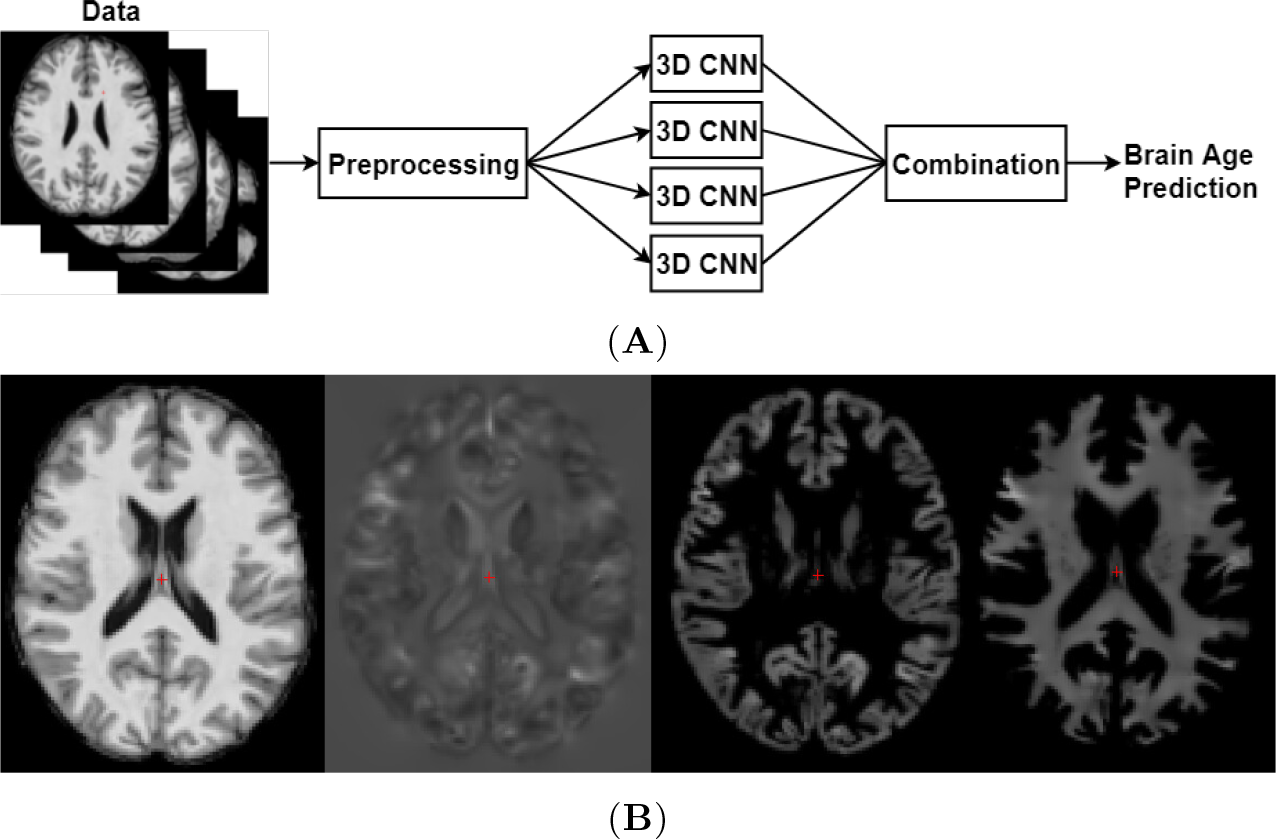
**(A)** A flowchart s howing a h igh-level o verview o f t he p roposed b rain a ge p rediction system. **(B)** Examples of image types generated by the preprocessing step. From left to right: a registered T1-weighted slice, a Jacobian map slice, a GM segmented slice and a WM segmented slice.

## 2. Results

### 2.1. Combining CNN outputs improves prediction accuracy

Our brain age prediction method was developed using images from structural brain MRIs for 1264 healthy Icelanders. To overcome problems caused by training a DL method on such a small dataset we use multiple images of the same individuals and utilize a data augmentation strategy. We start off by training the method independently on the four previously mentioned image types (Table 1A). The CNN that predicts the test set with the least error is the CNN trained on T1-weighted images followed by the CNN trained on WM segmented images.^1^

Having four predictions from four different data sources opens up the possibility of combining the predictions. The most straightforward way of fusing the forecasts is by using a majority voting scheme, *e.g.* by averaging the predictions made by the four CNNs. Another way to combine forecasts is to implement a data blender, for example, by implementing a linear regression model trained to predict brain age from the four CNN brain age predictions. This technique attempts to find the best linear combination of the four brain age predictions so in theory it should be guaranteed to be at least as good as the best predicting CNN method. To demonstrate this, we tried combining CNN brain age predictions using majority voting and linear regression data blending (Table 1B). Comparing the test set results of Table 1B to the results in Table 1A, we see that combining predictions results in lower test error than achieved by the CNN trained on T1-weighted images.

It is not straightforward to compare the accuracy of our method to previous brain age prediction methods, because they are evaluated on other datasets. However, to establish a baseline for the CNN based techniques, we investigated methods based on feature extraction such as surface-based morphometry (SBM) [19], voxel-based morphometry (VBM) [3], and similarity matrices. Machine learning regression methods were trained on these three types of features separately (Table 1C).^2^ If we compare the results in Table 1B and 1C we see that by combining CNN outputs we get predictions that are more accurate than those based on the feature extraction methods.

**Table 1:**
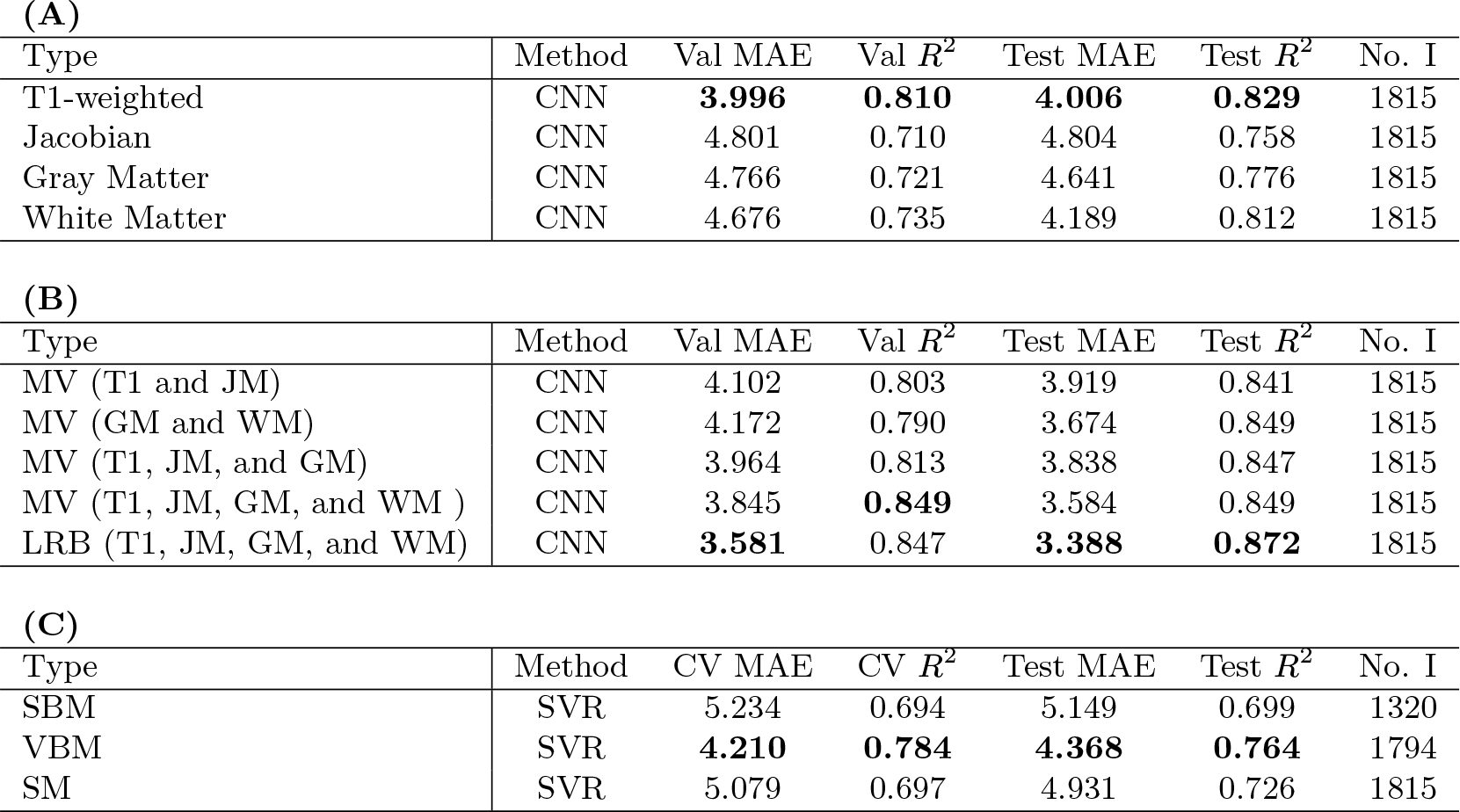
**(A)** The performance of the CNNs that were trained using T1-weighted images, Jacobian maps, GM and WM segmented images. Training set (*N* = 1171), validation set (*N* = 298), and test set (*N* = 346). The best results are shown in bold. **(B)** The performance when combining CNN predictions. The training/validation/test split is the same as for Table 1A. **(C)** The results of the best methods trained on SBM, VBM and similarity matrix features. The cross validation was performed using 10-fold cross validation. The SBM feature training/test split was 1056/264, the VBM feature training/test split was 1438/356, and the SM feature training/test split was 1469/346. Abbreviations: cross validation (CV), gray matter (GM), images (I), Jacobian map (JM), linear regression blender (LRB), majority voting (MV), mean absolute error (MAE), similarity matrix (SM), surface-based morphometry (SBM), validation (val), voxel-based morphometry (VBM), white matter (WM).

| **(A)** |  |  |  |  |  |  |
| --- | --- | --- | --- | --- | --- | --- |
| Type | Method | Val MAE | Val $R^2$ | Test MAE | Test $R^2$ | No. I |
| T1-weighted | CNN | **3.996** | **0.810** | **4.006** | **0.829** | 1815 |
| Jacobian | CNN | 4.801 | 0.710 | 4.804 | 0.758 | 1815 |
| Gray Matter | CNN | 4.766 | 0.721 | 4.641 | 0.776 | 1815 |
| White Matter | CNN | 4.676 | 0.735 | 4.189 | 0.812 | 1815 |

| **(B)** |  |  |  |  |  |  |
| --- | --- | --- | --- | --- | --- | --- |
| Type | Method | Val MAE | Val $R^2$ | Test MAE | Test $R^2$ | No. I |
| MV (T1 and JM) | CNN | 4.102 | 0.803 | 3.919 | 0.841 | 1815 |
| MV (GM and WM) | CNN | 4.172 | 0.790 | 3.674 | 0.849 | 1815 |
| MV (T1, JM, and GM) | CNN | 3.964 | 0.813 | 3.838 | 0.847 | 1815 |
| MV (T1, JM, GM, and WM ) | CNN | 3.845 | **0.849** | 3.584 | 0.849 | 1815 |
| LRB (T1, JM, GM, and WM) | CNN | **3.581** | 0.847 | **3.388** | **0.872** | 1815 |

| **(C)** |  |  |  |  |  |  |
| --- | --- | --- | --- | --- | --- | --- |
| Type | Method | CV MAE | CV $R^2$ | Test MAE | Test $R^2$ | No. I |
| SBM | SVR | 5.234 | 0.694 | 5.149 | 0.699 | 1320 |
| VBM | SVR | **4.210** | **0.784** | **4.368** | **0.764** | 1794 |
| SM | SVR | 5.079 | 0.697 | 4.931 | 0.726 | 1815 |

<sup>2</sup> Appendix B includes more information and results about of the regression methods trained on extracted features.

### 2.2. Testing the CNN on other datasets

Next, we examine how the method performs if we predict brain age of images from other datasets. To do so, we evaluate it on the IXI^3^ and UK Biobank [74] datasets and combine predictions using majority voting. We use this combination method rather than data blending, because it has similar accuracy to linear regression blender with the added benefit that it is unnecessary to train an extra linear model on the predictions. We observe that the initial prediction error of the method is high (Table 2). The problem is that there can be subtle differences between data from different scanning sites which will cause a model trained on one site to fail when predicting on the other site. There are multiple reasons for this. The MRI scanner type and parameters between sites can be different, which can cause differences between resolution, contrast and noise levels. Also, the distribution of age can be different between sites, for example, it is problematic if the new site has a wider age range than the training set.

We hypothesize that a CNN that is already proficient at predicting brain age at one site only needs a small adjustment to adapt to data from a new site. A transfer learning strategy achieves this: First, we freeze the model weights of the convolutional layers so that only the fully connected layers are trainable. Second, the CNN is re-trained on a portion of the data from the new site. An advantage of this strategy is that there are now fewer parameters to train, which means we can use less data and training will be faster. We carry out the transfer learning strategy by re-training the majority voting CNN on 440 images from the IXI dataset.

The re-trained CNN is validated on 104 images from the IXI dataset left out during training (validation set) and tested on 12395 images from the UK Biobank dataset (test set). Table 2 shows that the prediction accuracy is increased significantly by doing so.^4^ Surprisingly the accuracy of predictions for the UK Biobank site improve even though the CNN was not explicitly trained on it. This is intriguing and is perhaps explained by the fact that the IXI set includes a wider age range than the Icelandic set and includes 3T MRI images unlike the Icelandic set.

**Table 2:** UK Biobank and IXI prediction performance with and without transfer learning. The best results are shown in bold. Abbreviations: subjects (S), transfer learning (TL), validation (val).

| TL Used | IXI |  |  | UK Biobank |  |  |
| --- | --- | --- | --- | --- | --- | --- |
| | Val MAE | Val $R^2$ | Val Set Size | Test MAE | Test $R^2$ | Test Set Size |
| No | 6.420 | 0.778 | 104 | 8.494 | -0.630 | 12395 |
| Yes | **4.149** | **0.907** | 104 | **3.631** | **0.614** | 12395 |

### 2.3. Effect of random CNN weight initialization on PAD

We know that because CNNs start out in random initial states, and because they have highly non-convex loss functions [48], it is possible that two randomly initialized instances of our brain age prediction method will converge to two distinct local minima. These states could in theory both predict age equally well but have uncorrelated PAD values. Here we face a potential problem, because in the absence of a ground truth for the PAD there is no way to tell if either one of these PAD predictions is accurate. This sort of unreliable CNN behaviour would be problematic for any downstream analysis that utilizes the brain age prediction, because any conclusions made about the PAD would depend on the initialization of the CNN. In light of this, it would be reassuring if we could demonstrate that our method generally converges to similar PAD predictions after training.

To test this, four additional randomly initialized instances of our brain age prediction method are trained and the agreement between their PADs is examined. This procedure entails repeating these three main steps four times: **1)** Train four CNNs on the Icelandic dataset on the four previously mentioned image types. **2)** Freeze convolutions layers and train the CNNs on the IXI dataset (transfer learning step). **3)** Predict brain age in the UK Biobank dataset using CNNs, combine the predictions with majority voting and calculate PAD values.

After repeating these steps, we get four instances of the brain age prediction method that predict brain age of the 12395 subjects in the UK Biobank with mean absolute error (MAE) equal to 4.6, 5.5, 5.4, and 4.9 respectively. The reason why the error is higher here compared to the original results is that we did not spend as much time optimizing these CNNs. Nevertheless, if we look at the agreement of the original and the four new PADs we find that the intraclass correlation (ICC) is estimated to be equal to 0.86 (95% confidence interval [CI] = [0.855, 0.863]). This indicates that the UK Biobank PAD calculated using our method stays rather consistent between the five different training runs and is relatively robust to random weight initialization.

### 2.4. Associations between PAD and performance on neuropsychological tests

As mentioned above, previous studies have linked high PAD to cognitive impairment [21, 24, 10, 49]. In light of this, we are interested in looking at if PAD associates with performance on neuropsychological tests. Specifically, performance on tests administered by the UK Biobank that are designed to measure: fluid intelligence, numeric memory, visual memory, prospective memory, simple processing speed, complex processing speed, visual attention, and verbal fluency. To estimate PAD in the UK Biobank, we train four CNNs on the Icelandic set, then the IXI set using transfer learning, and combine their predictions using majority voting. We see from Table 3 that PAD is associated with worse performance on the digit substitution test (DSST), trail making tests (TMTs), and the reaction time test.^5^ As expected, these results indicate that PAD is in fact associated with cognitive impairment.

### 2.5. Genome-wide association study

PAD has previously been shown to be heritable [11, 35], however, to our knowledge no sequence variants conferring risk of or protecting against PAD have been identified. In order to look for such variants, we ran a genome wide association scan (GWAS) in the UK Biobank sample on PAD (same PAD as Section 2.4). This scan yields two sequence variants, rs2435204-G and rs1452628-T (Figure 2 and Table 4A). Additionally, given that sequence variants known to associate with brain structure are likely to be enriched for variants that associate with PAD. We decided to test a smaller set of 331 brain structure variants for association with PAD^6^. This yielded associations with three additional variants (Table 4B).

**Table 3:** Pearson’s r correlation between PAD and performance on neuropsychological tests. Negative DSST, positive TMT, and positive Reaction Time indicate worse performance. Abbreviations: confidence interval (CI), digit substitution test (DSST), trail making test (TMT).

| Neuropsychological Test | PAD Correlation (%) | 95% CI (%) | $P$ Value | No. Subjects |
| --- | --- | --- | --- | --- |
| DSST | -8.0 | (-10.4, -5.7) | 3.0e-11 | 6852 |
| TMT B | 7.9 | (5.4, 10.4) | 8.0e-10 | 6080 |
| TMT A | 5.4 | (2.9, 7.9) | 2.3e-05 | 6080 |
| TMT B - A | 5.1 | (2.6, 7.6) | 8.6e-05 | 5920 |
| Reaction Time | 3.3 | (1.6, 5.1) | 2.2e-04 | 12387 |

**Figure 2:**
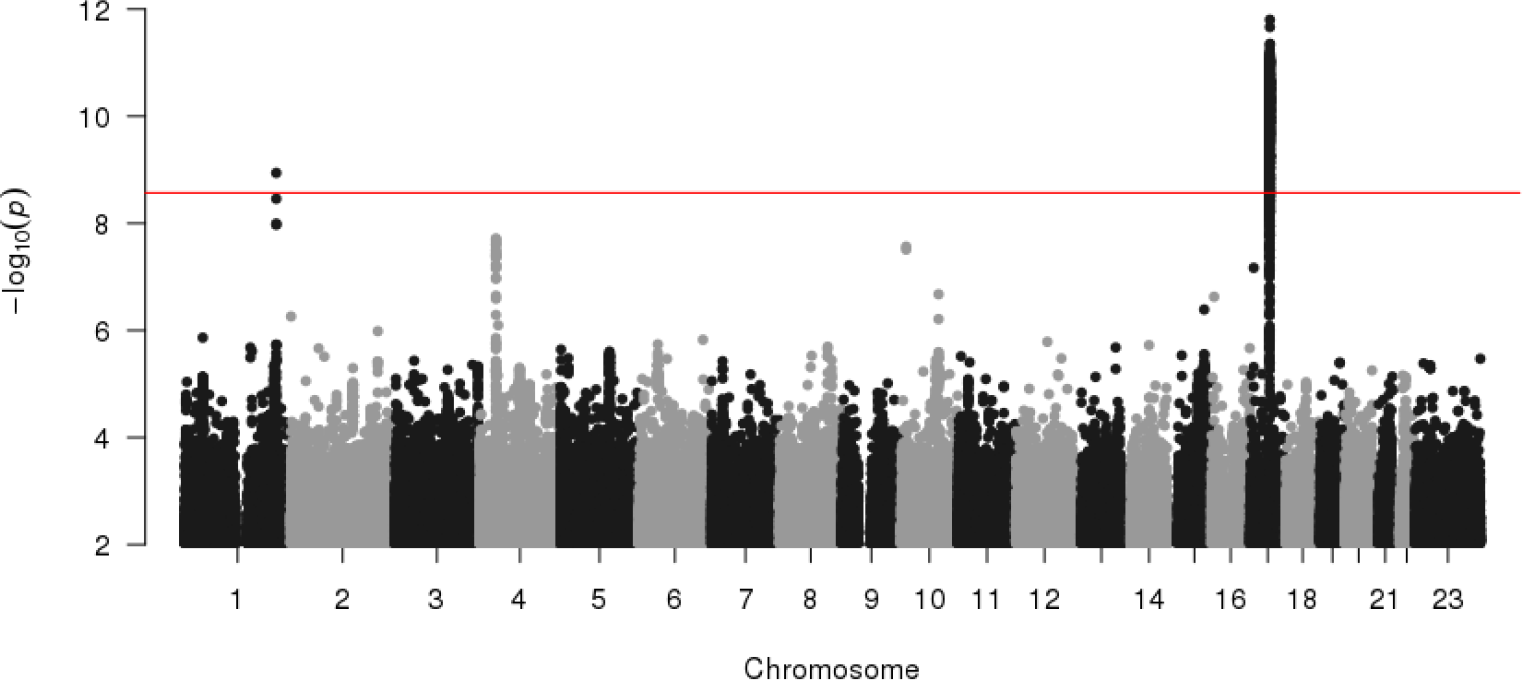
Manhattan plot of the GWAS results for the UK Biobank data. The horizontal line denotes the *P* value threshold for genome-wide significant effect.

The high number of tests conducted in GWAS combined with the general small effect size of common markers greatly increases the risk of a false postives [67]. To protect against potential confound effects we adjusted for variables, such as age, gender, total intracranial volume, and removed individuals of non-white British ancestry^7^. And then to thoroughly vet each hit we took three steps. **1)** We performed a replication test on held out data. **2)** Looked at whether the association results are affect by randomly initializing and retraining the CNN. **3)** Checked if the reported variants associate with other phenotypes related to brain ageing.

**Table 4:** Association between sequence variants and PAD for 12395 subjects. **(A)** Genome wide significant sequence variants. **(B)** Sequence variants associated with structural MRI brain phenotypes that also associate with PAD. Abbreviations: confidence interval (CI), minor allele frequency (MAF).

| **(A)** |  |  |  |  |  |  |
| --- | --- | --- | --- | --- | --- | --- |
| rs Number | Position<br>(GRCh38) | Allele<br>(min/maj) | MAF (%) | Effect | 95% CI | *P* Value |
| rs2435204 | chr17:45910839 | G/A | 26.6 | 0.10 | (0.07, 0.13) | 9.7e-12 |
| rs1452628 | chr1:214966544 | T/A | 36.2 | -0.08 | (-0.11, -0.05) | 1.2e-09 |

| **(B)** |  |  |  |  |  |  |
| --- | --- | --- | --- | --- | --- | --- |
| rs Number | Position<br>(GRCh38) | Allele<br>(min/maj) | MAF (%) | Effect | 95% CI | *P* Value |
| rs2790099 | chr6:45475612 | C/T | 36.0 | -0.06 | (-0.09, -0.03) | 5.8e-06 |
| rs6437412 | chr3:194776533 | C/T | 28.2 | -0.06 | (-0.09, -0.03) | 1.6e-05 |
| rs2184968 | chr6:126439848 | C/T | 46.0 | 0.05 | (0.03, 0.08) | 4.3e-05 |

**1)** The five reported sequence variants also associated with PAD in a replication set of 4453 subjects (Table 12 [Appendix G]). Four other variants which came up in the discovery stage were omitted because they did not replicate. **2)** The five sequence variants still associate with PAD even after we randomize the CNN weights and retrain the method (Table 13 [Appendix H]). This fits with the PAD ICC results from Section 2.3, which indicate that retraining the method should not have a large effect on any downstream analysis. The reason why the original PAD generally has a stronger association than the other four PAD estimates could be due to the Winner’s curse [80], or the fact that the new PAD estimates are less optimized. **3)** The identified sequence variants also associate with brain structure likely to be affected by brain ageing.^8^ Table 7 (Appendix F) shows that both PAD and rs1452628-T associate with lower cerebrospinal fluid (CSF) throughout the cerebral cortex which is consistent with reduced cortical sulcal openings. On the other hand, rs2435204-G associates with lower total white matter surface area, and reduced area in a number of cortical brain regions (Table 8). We also see that the other three sequence variants and PAD are associated with numerous structural brain phenotypes (Tables 9–11). In addition, we scanned for the phenotype effects of all five single-nucleotide polymorphisms (SNPs) in the UK Biobank data analyzed by the Roslin Institute [7] (Supplementary Tables 1-5).

## 3. Discussion

Here, we have presented a novel deep learning approach, using residual convolutional neural networks to predict brain age from a T1-weighted MRI, a Jacobian map, and gray and white matter segmented images, to study the discrepancy between age-related structural brain changes and chronological age. The MRI based deep learning system was shown to predict brain age from T1-weighted MRI data with a MAE = 3.39 and *R*^2^ = 0.87 on test data. Comparing our approach to other machine learning methods trained on surface-based morphometry, voxel-based morphometry, and similarity matrix features, we showed that our approach predicts brain age more accurately. We showed that transfer learning can be used to efficiently increase prediction accuracy for new sites. The PAD calculated using this method was shown to be relatively robust to random weight initialization and retraining, a result that indicates that the PAD estimated using our method can be used as a reliable phenotype in the study of brain ageing, as well as in the study of specific disorders of the brain. We also proposed that PAD could be an informative phenotype for genetic association studies, and indeed, our association analysis of PAD in a discovery set of 12395 subjects and replication set of 4453 subjects yielded five sequence variants.

The sequence variant with the strongest association, rs2435204-G, tags the H2 (inverted) form of the 17q21.31 inversion polymorphism [71]. This inversion spans approximately 1 Mb and includes 10 genes, including *MAPT*, a gene that encodes the tau protein which has been implicated in various dementias [58]. In addition, micro-deletions within the inversion are known to cause intellectual disability [40]. The H1 inversion haplotype has been associated with increased risk of Parkinson’s disease, male pattern baldness, and several other phenotypes, whereas H2 has been associated with a number of phenotypes including neuroticism [54], fibromyalgia [43], lower educational attainment, increased fecundity [39], and smaller intracranial volume^9^ (ICV) [33]. Due to the extensive linkage disequilibrium (LD) the 17q21.31 inversion region, reported markers for various associations in the region often differ between studies. For example, the most recent GWAS meta-analysis of Parkinson’s disease reports an association with rs17649553-T, that is fixated on and highly correlated with the H2-tagging rs2435204-G (*r*^2^ = 0.82, *D*^′^ = 1), with *OR* = 0.78 (95% *CI* = [0.76, 0.80]), *P* = 1.26 · 10^−68^ [15].

rs2435204-G also associates with brain structure phenotypes. Table 8 (Appendix F), shows that both PAD and rs2435204-G associate with increased thickness and decreased area in cortical brain regions. Interestingly, this pattern of increased thickness and decreased area has previously been associated with neuroticism [61]. Thus, lifestyle or phenotypes associated with a high neuroticism score, including anxiety, worry, fear, anger, frustration, depressed mood and loneliness may associate with PAD.

The other genome-wide significant sequence variant, rs1452628-T, is located close to *KCNK2* (also known as *TREK1*), which belongs to the two-pore domain potassium channel family and is mainly expressed in the brain [28]. In mice, *KCNK2* has been implicated in neuroinflammation [4], cerebral ischemia [6], and blood-brain barrier dysfunction [78]. rs1452628-T correlates with SNPs that have previously been associated with cortical sulcal opening and GM thickness, rs6667184 (*r*^2^ = 0.68), and rs864736 (*r*^2^ = 0.49) [44].

In addition, we identified three sequence variants associated with PAD by restricting the analysis to SNPs known a priori to associate with structural phenotypes. **1)** rs2790099-C is located in an intron of *RUNX2*, a gene that encodes the *RUNX2* protein which is essential for osteoblastic differentiation and skeletal morphogenesis and has been shown to play several roles in cell cycle regulation [73]. Supplementary Figure 12 shows that rs2790099-C is a possible cis-eQTL of *RUNX2* and it is most expressed in the basal ganglia (caudate and putamen). This lines up with the a priori brain structure GWAS that shows that rs2790099-C has genome-wide significant associations with white matter volume of regions in the basal ganglia (putamen and pallidum) (Table 9). **2)** rs6437412-C is an intron variant of *LINC01968* that associates with increased cortical CSF (Table 10) and hematological traits (Supplementary Table 4). **3)** rs2184968-C is located in an intron of *CENPW*, a gene that has previously been associated with traits, such as, height [55], cognitive performance [47], and male-pattern baldness [57]. Our analysis shows that rs2184968-C is associated with increased CSF in subcortical regions and increased size of the fourth ventricle (Table 11).

Confound effects are a problem for big imaging studies due to the huge number of imaging artifacts that can potentially influence both imaging and non-imaging variables of interest [67]. Some of the confound effects we have tried to control for are effects due to age, sex, head size, population structure, and scanner type. Head motion is another potentially problematic confound effect, because it causes reduction of estimated gray matter volume and thickness in MRI images similar to what we expect to see due to ageing [60]. While head motion is not important in the evaluation of our method (see Cole et al. [11]), it is potentially a problematic confound for GWAS analysis because certain clinical groups associate more with scanner motion. Elliott et al. [17] suggest to use fMRI-derived head motion estimates to correct for confound effects due to head motion when running GWAS analysis on brain structure phenotypes. We did try to correct PAD for head motion as they suggest, however, this correction only had a small effect on our results. Other potential confounds that we looked at were sample relatedness (the first 40 principal from components genetic ancestry analysis), genotyping array, and the assessment center were neuropsychological testing was performed. As with head motion, adjusting for these variables did not affect our results.

From our analysis we see that PAD associated with worse performance on neuropsychological tests, specifically poor performance on DSST, TMT, and the reaction time tests (Table 3). Interestingly, both the DSST and the reaction time test are designed to measure cognitive processing speed. The TMT is designed to asses visual attention. However, psychomotor speed is a factor in successful TMT performance [63]. Furthermore, a decline in processing speed along with impairment of reasoning, memory, and executive function are well documented to occur in age-associated cognitive decline [16]. As such, these results are in line with other studies that link high PAD to cognitive impairment [21, 24, 10, 49]. We note, that the association between PAD and TMT is consistent with the previous finding of Cole et al. [10]. However, the large dataset used here gives more conclusive results. Supporting this, we additionally find that schizophrenia, a brain disorder characterized by complex patterns of cognitive impairment, correlates with positive PAD (greater brain ageing than chronological age) and (Table 6 [Appendix E]).

In conclusion, we have presented a new method for predicting brain age using cutting-edge machine learning techniques. Our deep learning method produces a single measure (PAD) from raw MRI data that captures complex underlying correlated changes in MRI and can be used to study various traits and diseases, and in particular for genetic discovery. Using such a method represents one potential way for overcoming challenges with high dimensional data and multiple testing that plagues MRI research. By applying our method to large genomic datasets such as the UK Biobank has enabled us to identify novel genetic components that influence brain ageing. Further research into these components has potential to shed more light on the biological underpinnings of the ageing brain and its connection to various diseases and disorders.

## 4. Materials and methods

### 4.1. Datasets

The proposed method was evaluated on T1-weighted MR images from three independent datasets: an Icelandic dataset, the UK Biobank dataset, and the IXI dataset. DeCODE genetics provided the Icelandic MR data, consisting of scans from 1264 healthy subjects aged between 18 and 75 years. This dataset includes 1815 scans in total, since some subjects have several scans. The Icelandic data were acquired using two different scanners, a 1.5T Phillips Achieva scanner, and a 1.5T Siemens Magnetom Aera scanner. Scans were imaged using a T1-weighted gradient echo sequence (Philips Achieva: repetition time (TR) = 8.6 ms, echo time (TE) = 4.0 ms, flip angle (FA) = 8°, 170 slices, slice thickness = 1.2 mm, acquisition matrix = 192 × 192, field of view (FOV) = 240 × 240 mm; Siemens Aera: repetition time (TR) = 2400 ms, echo time (TE) = 3.54 ms, flip angle (FA) = 8°, 160 slices, slice thickness = 1.2 mm, acquisition matrix = 192 × 192, field of view (FOV) = 240 × 240 mm). Any serious neurological disorders were prescreened and removed. Additionally, we removed from the training and holdout sets subjects diagnosed with neurodevelopmental and mental disorders such as autism, bipolar disorder, intellectual disability, or schizophrenia, and subjects with any copy number variations previously associated with neurodevelopmental or psychiatric disorders.

The UK Biobank dataset^10^ consists of T1-weighted MR images of 15040 healthy subjects aged between 46 and 79 years old. The data were all collected using a 3T Siemens Skyra scanner. It is well-known that the presence of undetected population structure can lead to both false positive results and failure to detect genuine associations in genetic association studies [51], in an effort to combat this our analysis was constrained to 12395 individuals of white British ancestry. An additional release of MRI images by UK Biobank was added to a replication set. This set contains 6888 subjects (thereof 4458 subjects of white British ancestry) aged between 47 and 80 years old. The images in this set were collected using the same protocol as the previous UK Biobank set.

The IXI dataset consists of T1-weighted MR images of 544 healthy subjects and is freely available online. The subjects age at imaging was between 20 and 86 years old. The IXI data was collected from three different sites. The Hammersmith Hospital using a Philips 3T system, Guy’s Hospital using a Philips 1.5T system and the Institute of Psychiatry using a GE 1.5T system.^11^

### 4.2. Preprocessing

Preprocessing was carried out using the computational anatomy toolbox (CAT12) [23]. First, the input data were inhomogeneity corrected. Then the skull and other nonbrain elements were removed. Finally, the images were registered into the standard MNI space using the deformable registration algorithm DARTEL [2]. For further information, refer to the CAT12 manual [8].

There are three types of images that the preprocessing step generates. The first is an MNI-registered image. Second, a Jacobian map which is a by-product of the deformable registration. Lastly, a gray matter and white matter soft segmented images. All of the image types mentioned above have voxel size 1.5 mm^3^ and size 121×145×121.

### 4.3. CNN architecture

The CNN uses a residual architecture [26] as depicted in Figure 3. It consists of five residual blocks, each followed by a max pooling layer of stride 2×2×2 and kernel size 3×3×3, and one fully connected block. The convolutional part of the CNN reduces the input image from size 121×145×121 to 128 feature maps of size 4×5×4. The fully connected part reduces these feature maps down to an age prediction.

**Figure 3:**
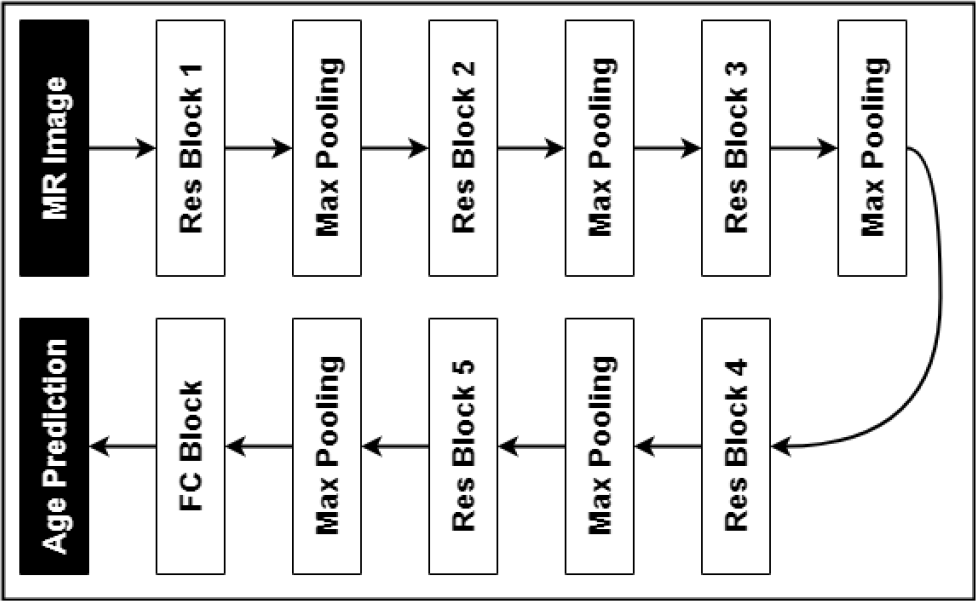
A flowchart showing the components of the proposed CNN architecture. Abbreviations: residual (Res), fully connected (FC).

The residual block, displayed in Figure 4, consists of a combination of layers which are repeated twice inside the residual blocks. This combination is composed of a 3D convolutional layer with stride 1×1×1 and kernel size 3×3×3, a batch re-normalization layer [34], and an ELU activation function [9]. The defining element of the residual block is the skip connection which adds the signal feeding into the residual block to the output of a layer close to the end of the block. The number of feature maps in block number *n* was chosen by the rule 2^*n*+2^.

**Figure 4:**
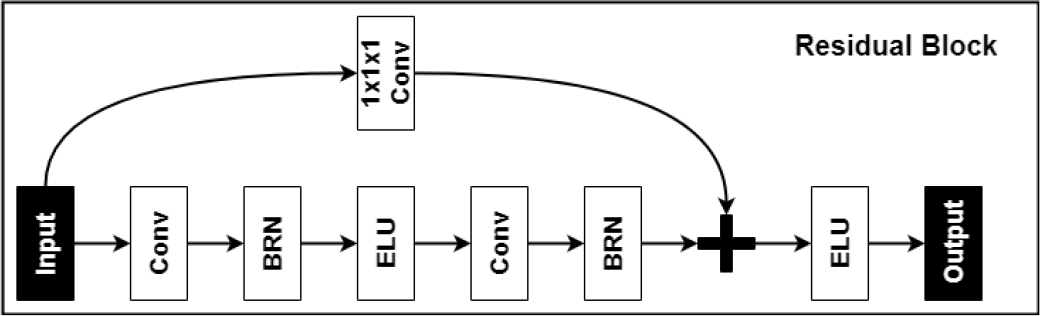
A flowchart showing the components of the proposed residual block. Abbreviations: batch re-normalization (BRN), convolutional layer (Conv).

The fully connected block, depicted in Figure 5, is a multilayer perceptron (MLP) [81] with one hidden layer. The input layer has 128 × 4 × 5 × 4 = 10240 neurons, the hidden layer (FC 1) has 256 neurons that use an ELU activation function, and the output layer has a single neuron. Following the hidden layer, a dropout [69] layer with keep rate equal to 0.8 is employed. The output layer (FC 2) has no activation function which means that it performs a linear regression on the hidden layer features. To account for factors such as scanner type and sex that can affect the estimated brain age of an individual we include them as inputs in the linear regression by concatenating them with the hidden features of the MLP.

**Figure 5:**
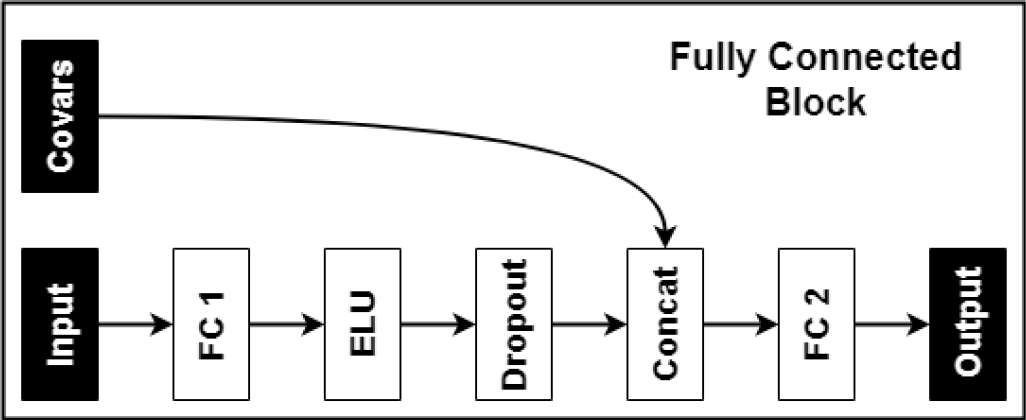
A flowchart showing the components of the proposed fully connected block. Abbreviations: fully connected layer one (FC1), concatenation layer (Concat), fully connected layer two (FC2).

The mean absolute error was used as the loss function and the CNN was optimized using Adam [37] with parameters: learning rate = 0.001, decay= 10^−6^, *β*_1_ = 0.9, *β*_2_ = 0.999, and batch size = 4. The He initialization strategy [27] was used to initialize the weights, and each trainable node in the CNN was regularized with *l*_2_ weight decay [42], with *λ* = 5 × 10^−5^. Early stopping [53] was used, i.e., if the validation error did not improve in 100 epochs the training was stopped. Furthermore, to reduce the risk of overfitting, data augmentation [25] was used to generate new training instances by applying a coordinate transformation to a random subset of the training data, consisting of a combined 3D rotation and a 3D translation. The rotation angles were between −40 and 40 degrees with equal probability, and the translation distance, for each direction, was selected between −10 and 10 voxels with equal probability.

### 4.4. SBM, VBM, and similarity matrix brain age prediction

The SBM features were generated using FreeSurfer’s recon-all algorithm^12^ [18] and the VBM features were generated using the CAT12 toolbox^13^. The similarity matrix was constructed by taking the inner product between the combined gray and white matter segmented images of each subject. The SBM and VBM features were adjusted for intracranial volume, sex and scanner type. The features were then zero centered and normalized to unit variance. The regression methods that were tested were, linear regression [65], lasso [75], ridge regression [30], elastic net [82], random forest regression [29], and SVR [68]. A grid search was used to find the tuning parameters corresponding to the lowest cross-validation error for the methods mentioned.

### 4.5. Statistical methods

To assessing the accuracy of the machine learning methods we performed simple training and validation splits, and selected a suitable model by evaluating the validation MAE. The subjects from the Icelandic sample were split between these three sets, and if a subject had multiple images, the images were all put in the same set. The data were divided into 64% training set (*N_s_* = 809, *N_i_* = 1171), 16% validation set (*N_s_* = 202, *N_i_* = 298), and 20% test set (*N_s_* = 253, *N_i_* = 346), were *N_s_* is the number of subjects and *N_i_* is the number of images. When evaluation the machine learning models the MAE and *R*^2^ score for the images in the validation and test set is calculated.

To assess the transfer learning performance, the IXI dataset was split into 80% training set (*N* = 440), 20% validation set (*N* = 104) and the whole UK Biobank dataset was used as a test set (*N* = 12395). As before, we evaluate accuracy by calculating the MAE and *R*^2^ score on the validation and test set.

In order to test the reliability of PAD, the intraclass correlation was calculated with ICCbare from the ICC R package^14^. The 95% confidence interval was estimated using bootstrapping with 2000 sampling iterations.

The Pearson correlation coefficient was calculated in order to test for association between PAD and performance on neuropsychological tests. Before performing the analysis we first adjusted the PAD for age, age^2^, total intracranial volume, sex, the interaction between sex and age. The correction was performed using linear regression. Ten correlation tests were performed, so a Bonferroni adjusted significance level *α_B_*_2_ = 0.05/10 = 0.005. We performed a GWAS on PAD to find associated sequence variants. For the genetic analysis we used version 3 of the imputed genetic dataset released by UK Biobank in July 2017 [5]. The UK Biobank genetic data was assayed using two very similar genotyping arrays (95% of marker content is shared). Roughly 10% of the subjects were genotyped using applied Biosystems UK BiLEVE Axiom Array by Affymetrix and the rest using the closely related Applied Biosystems UK Biobank Axiom Array [5]. Variants with imputation quality score below 0.3, and minor allele frequency below 0.1% were filtered out, which left ~20 million variants to be considered for GWAS. Before performing GWAS, the PAD was adjusted for age, age^2^, total intracranial volume, sex, the interaction between sex and age using linear regression. The adjusted PAD was then normalized with an inverse normal transformation. Sequence variants associated with PAD are only reported if they reach genome-wide significance. If two genome-wide significant variants are in LD (*r*^2^ > 0.1) we report the variant with the lower p-value.

In addition, we tested for association between PAD and sequence variants known to associate with structural brain phenotypes. These variants were found by performing GWAS separately on 305 SBM phenotypes generated with recon-all by using the Freesurfer 6.0 software [18] and 540 VBM phenotypes generated by using CAT12 [23]. All genome-wide significant markers were then aggregated into a single list. In cases where variants were in LD (*r*^2^ > 0.5), only the variant with the lower p-value was selected. The final list included 331 variants, to account for testing test variants for the second time a Bonferroni adjusted significance level 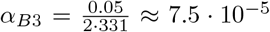 was used for the PAD association test.

To reduce the risk of false positive sequence variant associations we additionally checked for association in a replication set of 4453 subjects. To pass this test the association between the variants under consideration and PAD need to show evidence of statistical significance (*α_R_* < 0.05).

### 4.6. eQTL analysis

To investigate if any of the variants are expression quantitative trait loci (eQTLs) we used the GTEx database^15^ [13]. Our eQTL analysis was carried out by logging onto https://gtexportal.org, typing in the corresponding rs number of identified variants, and checking if they have any associated eQTLs. However, identifying whether a variant is truly causal in both GWAS and eQTL is challenging because of the uncertainty caused by LD [31]. Therefore, we only report variants as eQTLs of genes if they are close to being the most significant eQTL of that specific gene.

## Code availability

Any custom code or software used to implement the brain age prediction method detailed in this paper will be made available upon request.

## Data availability

The genetic and phenotype datasets generated by UK Biobank used in this study are available via the UK Biobank data access process (see http://www.ukbiobank.ac.uk/register-apply/). Detailed information about the genetic data and MRI data available in UK Biobank is listed here: http://www.ukbiobank.ac.uk/scientists-3/genetic-data/, https://www.fmrib.ox.ac.uk/ukbiobank/.

The Icelandic data used in this publication are not publicly available due to information, contained within them, that could compromise research participant privacy. The authors declare that the data supporting the findings of this study are available within the article, its supplementary information, and upon request.

## Acknowledgment

This research has been conducted using the UK Biobank Resource under Application Number 24898. The research leading to these results has received support from the Innovative Medicines Initiative Joint Undertaking under grant agreements no. 115008 (NEWMEDS) and no. 115300 (EUAIMS), of which resources are composed of EFPIA in-kind contribution and financial contribution from the European Union’s Seventh Framework Programme (EU-FP7/2007-2013). The financial support from the European Commission to the NeuroPain project (FP7#HEALTH-2013-602891-2) is acknowledged.

## Author contributions

B.A.J. implemented the method, wrote the code, and performed experiments. B.A.J and M.O.U. developed the method and designed statistical experiments. All authors contributed to the final version of the manuscript.

## Additional results

## Appendix A: Chronological age distribution of the datasets

Figures 6, 7, 8 show histograms of the distribution of age in the Icelandic dataset, UK Biobank dataset, and IXI dataset.

**Figure 6:**
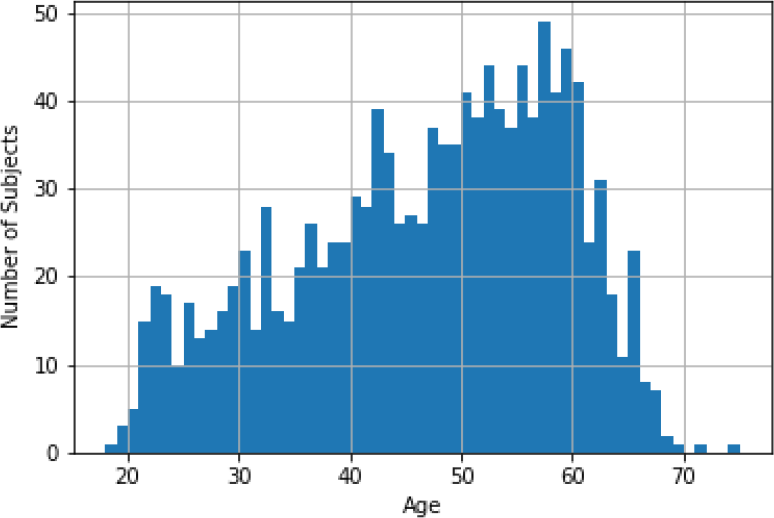
A histogram showing the distribution of chronological age in the Icelandic dataset.

**Figure 7:**
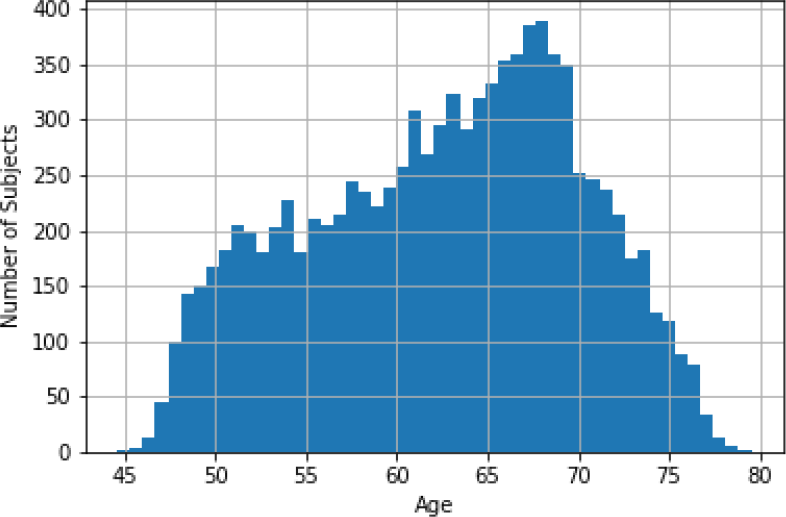
A histogram showing the distribution of chronological age in the UK Biobank dataset.

**Figure 8:**
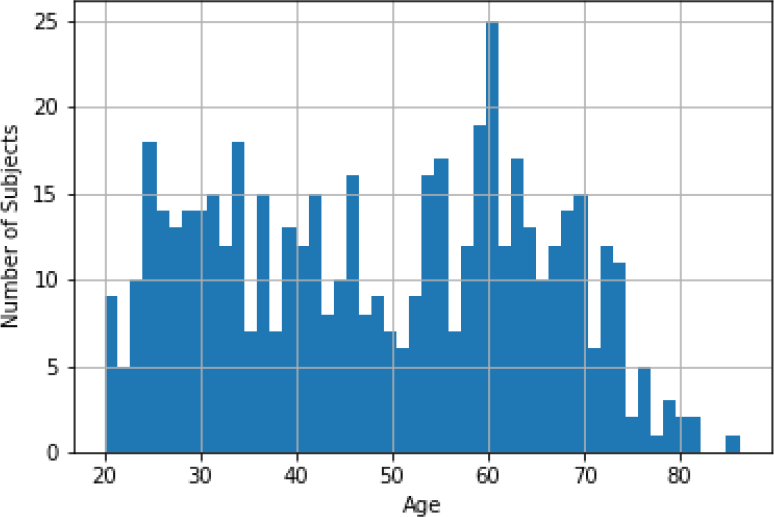
A histogram showing the distribution of chronological age in the IXI dataset.

## Appendix B: SBM, VBM, and similarity matrix results

All of the SBM, VBM, and similarity matrix results are shown in Table 5. The method that performed best on the SBM features was a radial basis function (RBF) kernel SVR method with *C* = 10 and *γ* = 10^−3^. Similarly, the best performing method on VBM features was an RBF kernel SVR method with parameters *C* = 100 and *γ* = 10^−3^. The best performing method on similarity matrix features was a linear kernel SVR method with penalty parameter *C* = 100.

**Table 5:** All SBM, VBM, and similarity matrix results. The cross validation was performed using 10-fold cross validation. Abbreviations: cross validation (CV), linear regression (LR), ridge regression (RR), elastic net (EN), random forest (RF), similarity matrix (SM), subjects (S), images (I).

| Method | Type | CV MAE | CV $R^2$ | Test MAE | Test $R^2$ | No. S | No. I |
| --- | --- | --- | --- | --- | --- | --- | --- |
| LR | SBM | 6.218 | -1.516 | 5.239 | 0.677 | 1263 | 1320 |
| RR | SBM | 5.275 | 0.688 | 5.149 | 0.699 | 1263 | 1320 |
| Lasso | SBM | 5.359 | 0.679 | 5.324 | 0.685 | 1263 | 1320 |
| EN | SBM | 5.276 | 0.689 | 5.149 | 0.699 | 1263 | 1320 |
| RF | SBM | 6.069 | 0.588 | 6.365 | 0.557 | 1263 | 1320 |
| SVR | SBM | 5.234 | 0.694 | 5.149 | 0.699 | 1263 | 1320 |
| LR | VBM | 4.769 | 0.729 | 5.034 | 0.699 | 1246 | 1794 |
| RR | VBM | 4.579 | 0.753 | 4.907 | 0.717 | 1246 | 1794 |
| Lasso | VBM | 4.596 | 0.756 | 4.805 | 0.724 | 1246 | 1794 |
| EN | VBM | 4.545 | 0.759 | 4.859 | 0.719 | 1246 | 1794 |
| RF | VBM | 5.374 | 0.664 | 5.272 | 0.645 | 1246 | 1794 |
| SVR | VBM | 4.210 | 0.784 | 4.368 | 0.764 | 1246 | 1794 |
| LR | SM | 8.363 | 0.144 | 8.478 | 0.146 | 1264 | 1815 |
| RR | SM | 4.823 | 0.729 | 4.937 | 0.728 | 1264 | 1815 |
| Lasso | SM | 5.160 | 0.695 | 5.306 | 0.688 | 1264 | 1815 |
| EN | SM | 5.050 | 0.702 | 5.206 | 0.701 | 1264 | 1815 |
| RF | SM | 7.910 | 0.277 | 7.520 | 0.342 | 1264 | 1815 |
| SVR | SM | 5.079 | 0.697 | 4.931 | 0.726 | 1264 | 1815 |

## Appendix C: The neuropsychological tests

*Fluid intelligence*^16^: Participants are asked to solve problems that require logic and reasoning ability, independent of acquired knowledge and have 2 minutes to complete as many questions as possible.

*Numeric memory*^17^: Participants are shown a 2-digit number which then disappears and after certain period are asked to recall the number. The test starts with a 2-digit number and becomes 1-digit longer each time they remember correctly up to a maximum of 12 digits.

*Visual memory*^18^: Participants are asked to memorize the position of as many matching pairs of cards as possible. The cards are then turned face down and the participant is asked to find as many pairs as possible.

*Prospective memory*^19^: Participants are shown four colored shapes and asked to touch a square. They should remember that earlier they were asked to touch the orange circle instead.

*Simple processing speed*^20^: Participants play 12 rounds of the card-game ‘Snap’ to assess reaction time. They are shown two cards at a time; if both cards are the same, they press a button as quickly as possible.

*Complex processing speed*^21^: Participants are asked to solve a digit symbol substitution test (DSST). They are presented with a series of grids in which symbols are to be matched to numbers according to a key presented on the screen.

*Visual attention* ^22^: Participants are asked to solve a trail making test (TMT) of type A and B. They are presented with a series of labeled circles and instructed to touch them according to a particular ordering rule.

*Verbal fluency*^23^: Participants are interviewed by trained staff to assess cognitive function, based on how many words beginning with the letter ‘S’ they can state within one minute. This test is only available for a small subset of the total participants.

## Appendix D: Brain age prediction scatter plots

**Figure 9:**
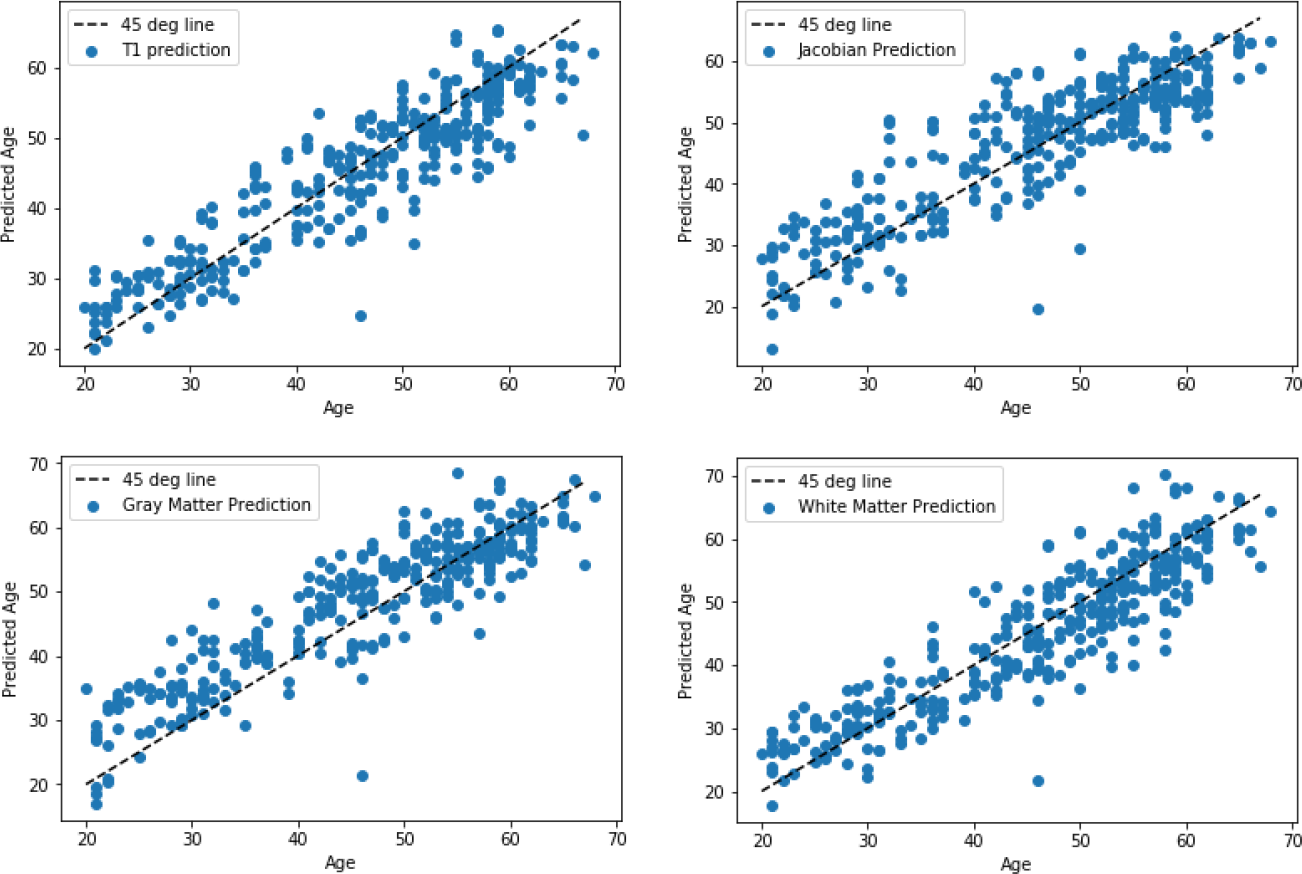
Scatter plots showing test set predictions made by CNNs. These plots show the chronological age against the brain age predicted by the CNNs trained on T1-weighted images, Jacobian maps, gray matter segmented images, and white matter segmented images. The top left plot shows the predictions made by the CNN trained on T1-weighted images. The top right plot shows the predictions made by the CNN trained on Jacobian maps. The bottom left plot shows the predictions made by the CNN trained on segmented gray matter images. The bottom right plot shows the predictions of the CNN trained on segmented white matter images.

**Figure 10:**
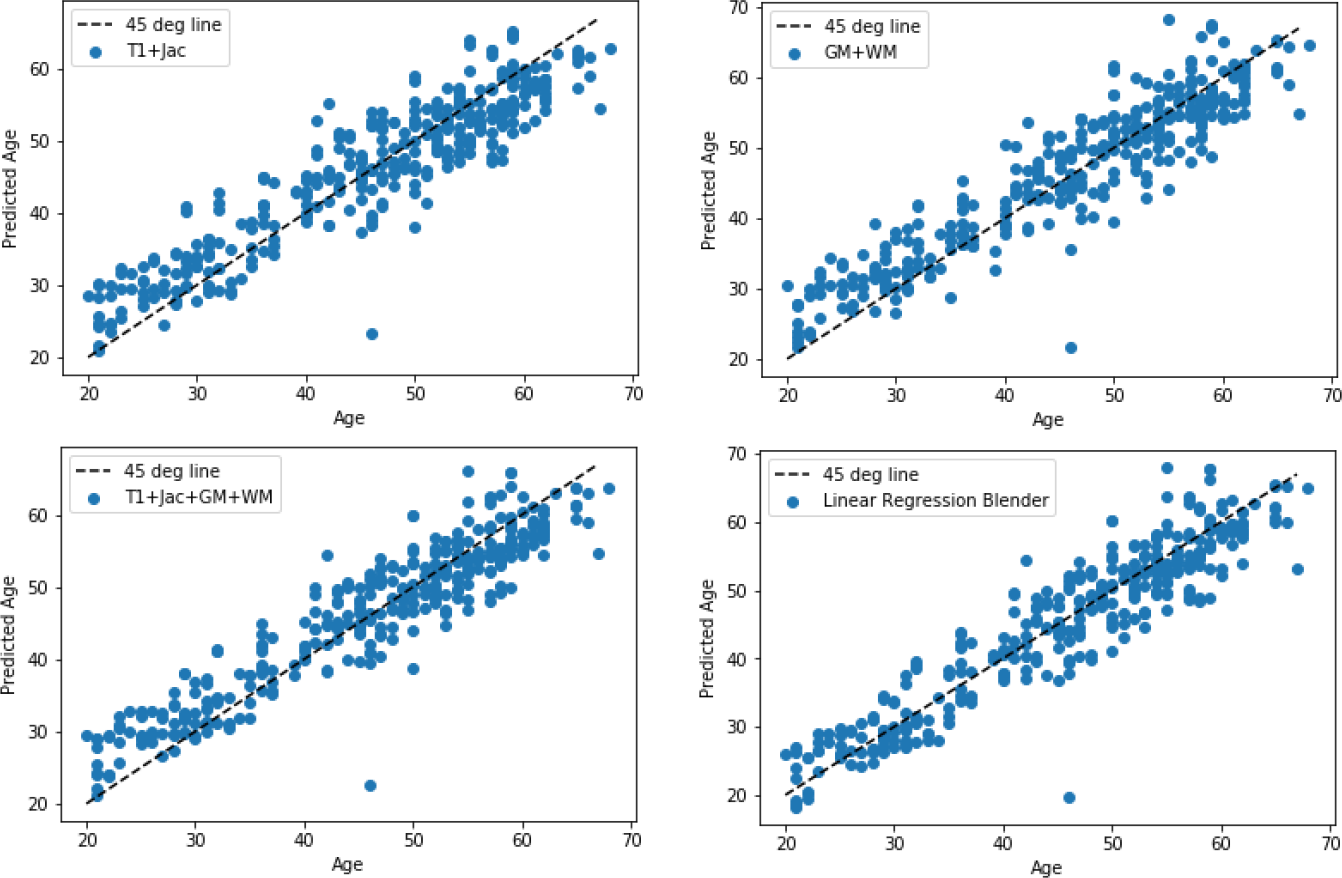
Scatter plots showing test set predictions made by combined CNN predictions. These plots show the chronological age against the brain age predicted by a combination of CNNs trained on registered T1-weighted images, Jacobian maps, gray matter segmented images, and white matter segmented images. The top left plot shows majority voting predictions made by the CNN trained on T1-weighted images and Jacobian maps. The top right plot shows the majority voting predictions made by CNNs trained on gray and white matter segmented images. The bottom left plot shows the majority voting predictions made by CNNs trained on T1-weighted images, Jacobian maps, segmented gray and white matter images. The bottom right plot shows the predictions made by the linear regression blender, trained on four predictions from CNNs trained on T1-weighted images, Jacobian maps, and segmented gray and white matter images.

**Figure 11:**
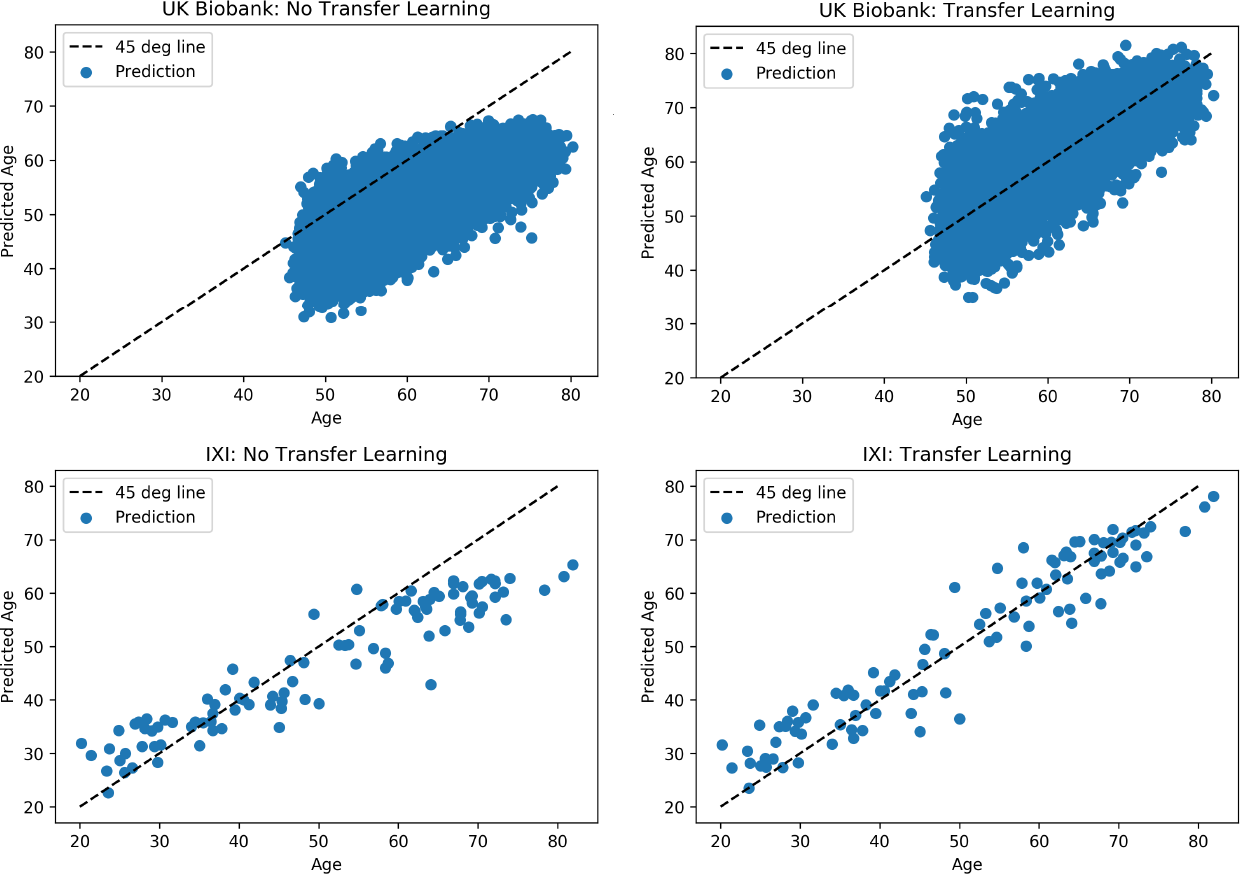
Four scatter plots that show the effect of using transfer learning, when predicting brain age of the IXI (validation set) and UK Biobank (test set) datasets, using a CNN trained on images from the Icelandic dataset. These plots show the chronological age against brain age predicted by the CNNs trained on T1-weighted images. The top left plot shows the UK Biobank brain age predictions without transfer learning. The top right plot shows the UK Biobank brain age predictions with transfer learning. The bottom left plot shows the IXI brain age predictions without transfer learning. The bottom right plot shows the IXI brain age predictions with transfer learning.

## 4.7. Appendix E: Predicting brain age of neurodevelopmental and mental disorder cases

Subjects from the Icelandic sample diagnosed with neurodevelopmental or mental disorders were left out while training the brain age prediction method. This includes subjects with autism, bipolar disorder, schizophrenia, and intellectual disability. We now look for deviation in PAD in these subjects compared to the PAD of the healthy Icelanders.

A two sample t-test was used to test for the difference between the mean PAD of the neurodevelopmental and mental disorder cases and healthy controls. Before performing the analysis we first compute the PAD. In cases where subjects had multiple images the average PAD was used instead. In this particular instance the PAD was adjusted for age, gender, and total intracranial volume using a generalized additive model. Five correlation tests were performed, so we used a Bonferroni adjusted significance level *α*_*B*1_ = 0.05/5 ≈ 0.01. The controls are taken from the Icelandic test set (N=291) and their average PAD is 0.1. Most of the subjects in the schizophrenia group are young males. Therefore, in an effort to compare similar groups we performed an additional test where only the PAD of males under 35 years is compared. After applying this constraint there are 22 control subjects left and their average PAD is −0.6.

**Table 6:** The average PAD difference between neurodevelopmental/mental disorder cases and controls from the test set. Abbreviations: confidence interval (CI).

| Cases | PAD Difference | 95% CI | P Value | No. Cases | No. Controls |
| --- | --- | --- | --- | --- | --- |
| Schizophrenia | 2.2 | (1.2, 3.2) | 3.5e-05 | 68 | 291 |
| Schizophrenia (Male Under 35) | 3.2 | (1.4, 5.1) | 1.1e-03 | 48 | 22 |
| Intellectual Disability | 1.5 | (-0.5, 3.6) | 1.2e-01 | 6 | 291 |
| Autism | 2.3 | (-0.9, 5.6) | 1.4e-01 | 10 | 291 |
| Bipolar Disorder | 0.4 | (-0.6, 1.5) | 4.0e-01 | 31 | 291 |

Table 6 shows that PAD is higher in individuals with schizophrenia than in controls. This is consistent with findings from other studies [56, 64, 41, 35] that have looked at brain ageing of schizophrenia patients. Brain structure irregularities in schizophrenia, such as cortical thinning [76] and cerebral ventricular enlargement [79] also seen in healthy ageing [62, 36] may be driving the prediction. Why there are structural differences in brains from schizophrenia patients is not fully understood. Psychotic episodes and longtime use of antipsychotic drugs [52] probably contribute. Sequence variants conferring high-risk of the disease have also been shown to affect brain structure in controls [70].

## Appendix F: SBM and VBM phenotypes associated with variants from brain age GWAS

**Table 7:**
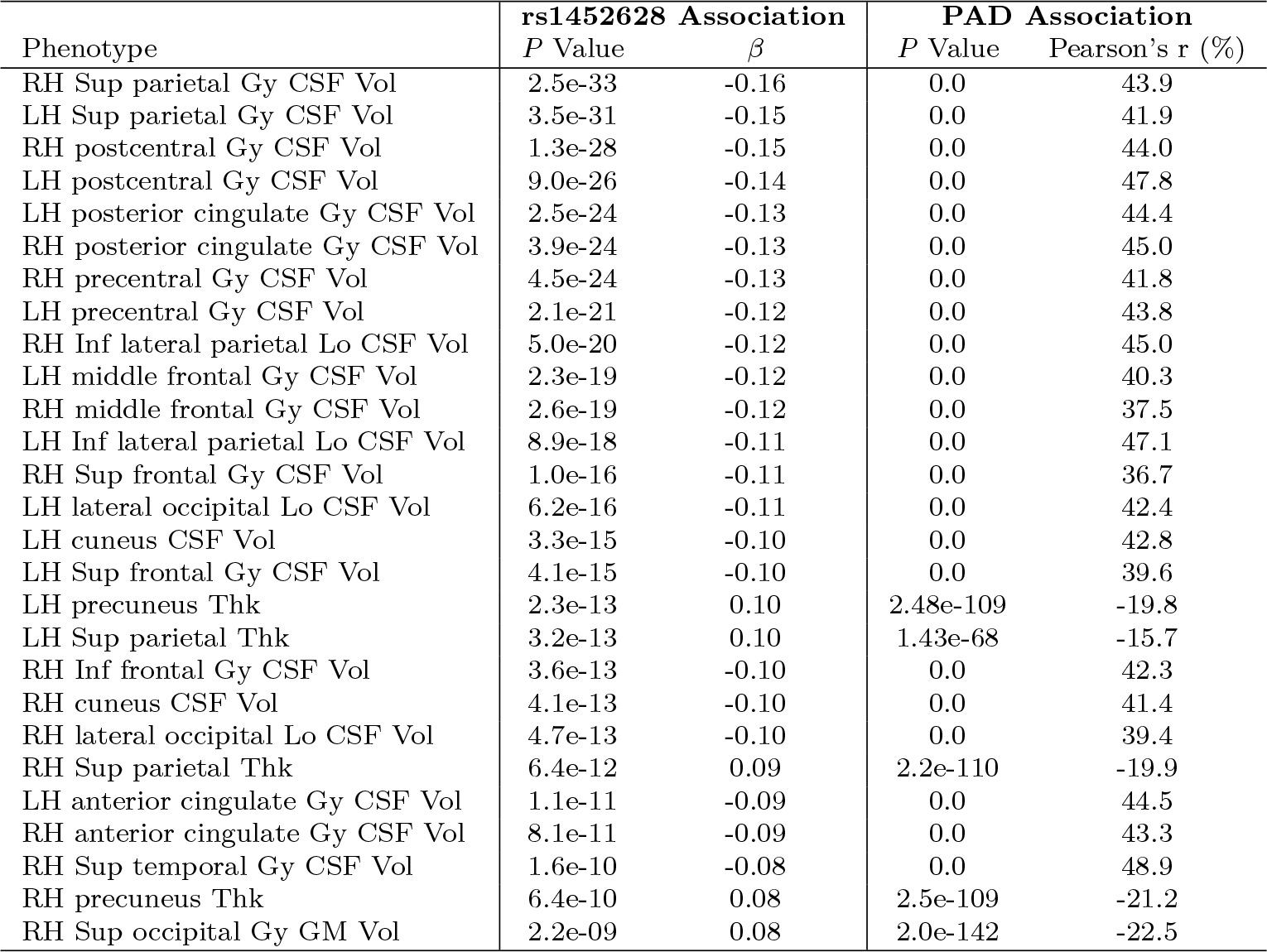
SBM and VBM phenotypes associated rs1452628. Only structural brain phenotypes that reach genome-wide significance in the discovery set (UK Biobank) are listed. In addition, the correlations between PAD and the SBM/VBM phenotypes are shown. Abbreviations: cerebrospinal fluid (CSF), gray matter (GM), gyrus (Gy), inferior (Inf), left hemisphere (LH), lobe (Lo), right hemisphere (RH), superior (Sup), thickness (Thk), volume (Vol).

| Phenotype | rs1452628 Association |  | PAD Association |  |
| --- | --- | --- | --- | --- |
| | *P* Value | $\beta$ | *P* Value | Pearson's *r* (%) |
| RH Sup parietal Gy CSF Vol | 2.5e-33 | -0.16 | 0.0 | 43.9 |
| LH Sup parietal Gy CSF Vol | 3.5e-31 | -0.15 | 0.0 | 41.9 |
| RH postcentral Gy CSF Vol | 1.3e-28 | -0.15 | 0.0 | 44.0 |
| LH postcentral Gy CSF Vol | 9.0e-26 | -0.14 | 0.0 | 47.8 |
| LH posterior cingulate Gy CSF Vol | 2.5e-24 | -0.13 | 0.0 | 44.4 |
| RH posterior cingulate Gy CSF Vol | 3.9e-24 | -0.13 | 0.0 | 45.0 |
| RH precentral Gy CSF Vol | 4.5e-24 | -0.13 | 0.0 | 41.8 |
| LH precentral Gy CSF Vol | 2.1e-21 | -0.12 | 0.0 | 43.8 |
| RH Inf lateral parietal Lo CSF Vol | 5.0e-20 | -0.12 | 0.0 | 45.0 |
| LH middle frontal Gy CSF Vol | 2.3e-19 | -0.12 | 0.0 | 40.3 |
| RH middle frontal Gy CSF Vol | 2.6e-19 | -0.12 | 0.0 | 37.5 |
| LH Inf lateral parietal Lo CSF Vol | 8.9e-18 | -0.11 | 0.0 | 47.1 |
| RH Sup frontal Gy CSF Vol | 1.0e-16 | -0.11 | 0.0 | 36.7 |
| LH lateral occipital Lo CSF Vol | 6.2e-16 | -0.11 | 0.0 | 42.4 |
| LH cuneus CSF Vol | 3.3e-15 | -0.10 | 0.0 | 42.8 |
| LH Sup frontal Gy CSF Vol | 4.1e-15 | -0.10 | 0.0 | 39.6 |
| LH precuneus Thk | 2.3e-13 | 0.10 | 2.48e-109 | -19.8 |
| LH Sup parietal Thk | 3.2e-13 | 0.10 | 1.43e-68 | -15.7 |
| RH Inf frontal Gy CSF Vol | 3.6e-13 | -0.10 | 0.0 | 42.3 |
| RH cuneus CSF Vol | 4.1e-13 | -0.10 | 0.0 | 41.4 |
| RH lateral occipital Lo CSF Vol | 4.7e-13 | -0.10 | 0.0 | 39.4 |
| RH Sup parietal Thk | 6.4e-12 | 0.09 | 2.2e-110 | -19.9 |
| LH anterior cingulate Gy CSF Vol | 1.1e-11 | -0.09 | 0.0 | 44.5 |
| RH anterior cingulate Gy CSF Vol | 8.1e-11 | -0.09 | 0.0 | 43.3 |
| RH Sup temporal Gy CSF Vol | 1.6e-10 | -0.08 | 0.0 | 48.9 |
| RH precuneus Thk | 6.4e-10 | 0.08 | 2.5e-109 | -21.2 |
| RH Sup occipital Gy GM Vol | 2.2e-09 | 0.08 | 2.0e-142 | -22.5 |

**Table 8:** SBM and VBM phenotypes associated with rs2435204. Only structural brain phenotypes that reach genome-wide significance in the discovery set (UK Biobank) are listed. In addition, the correlations between PAD and the SBM/VBM phenotypes are shown. Abbreviations: left hemisphere (LH), right hemisphere (RH), thickness (Thk), volume (Vol), white matter (WM).

| Phenotype | rs2435204 Association |  | PAD Association |  |
| --- | --- | --- | --- | --- |
| | *P* Value | $\beta$ | *P* Value | Pearson's *r* (%) |
| Total RH WM surface area | 5.1e-17 | -0.13 | 6.4e-27 | -9.67 |
| Total LH WM surface area | 4.5e-16 | -0.12 | 2.3e-30 | -10.3 |
| RH fusiform area | 1.8e-15 | -0.12 | 1.5e-22 | -8.80 |
| LH fusiform area | 3.1e-14 | -0.11 | 1.4e-20 | -8.38 |
| RH lateral occipital area | 8.9e-12 | -0.10 | 2.1e-7 | -4.68 |
| LH postcentral area | 2.2e-10 | -0.10 | 5.0e-2 | -1.77 |
| RH rostral middle frontal Thk | 2.5e-10 | 0.10 | 3.3e-52 | -13.7 |
| LH lateral occipital area | 9.8e-10 | -0.09 | 5.6e-8 | -4.90 |
| RH lingual area | 1.5e-09 | -0.09 | 8.7e-5 | -3.54 |
| Total LH cerebral WM Vol | 1.8e-09 | -0.09 | 5.0e-98 | -18.8 |

## Appendix F: SBM and VBM phenotype associations for structural brain prior variants

**Table 9:**
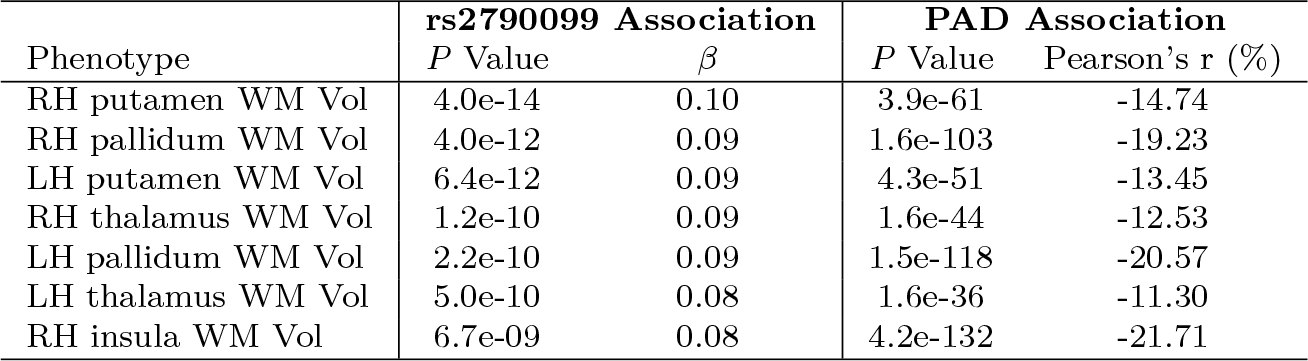
SBM and VBM phenotypes associated with rs2790099. Only structural brain phenotypes that reach genome-wide significance in the discovery set (UK Biobank) are listed. In addition, the correlations between PAD and the SBM/VBM phenotypes are shown. Abbreviations: left hemisphere (LH), right hemisphere (RH), volume (Vol), white matter (WM).

**Table 10:**
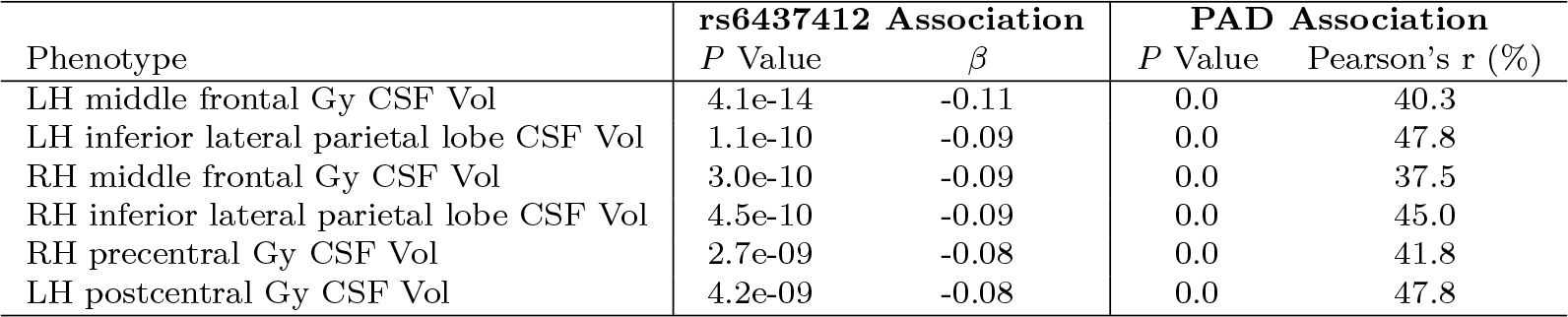
SBM and VBM phenotypes associated with rs6437412. Only structural brain phenotypes that reach genome-wide significance in the discovery set (UK Biobank) are listed. In addition, the correlations between PAD and the SBM/VBM phenotypes are shown. Abbreviations: cerebrospinal fluid (CSF), left hemisphere (LH), right hemisphere (RH), volume (Vol).

**Table 11:** SBM and VBM phenotypes associated with rs2184968. Only structural brain phenotypes that reach genome-wide significance in the discovery set (UK Biobank) are listed. In addition, the correlations between PAD and the SBM/VBM phenotypes are shown. Abbreviations: amygdaloid body (AmB), cerebrospinal fluid (CSF), gyrus (Gy), left hemisphere (LH), parahippocampal (ParHip), right hemisphere (RH), volume (Vol).

| Phenotype | **rs2184968 Association** |  | **PAD Association** |  |
| --- | --- | --- | --- | --- |
| | *P* Value | $\beta$ | *P* Value | Pearson's *r* (%) |
| RH cerebellum CSF Vol | 4.3e-20 | 0.12 | 0.0 | 39.7 |
| LH cerebellum CSF Vol | 1.4e-19 | 0.12 | 0.0 | 39.2 |
| RH brain stem CSF Vol | 3.1e-16 | 0.10 | 0.0 | 37.0 |
| LH brain stem CSF Vol | 3.9e-13 | 0.10 | 0.0 | 37.0 |
| LH AmB and ParHip Gy CSF Vol | 1.1e-09 | 0.08 | 3.6e-316 | 33.2 |
| 4th Ventricle Vol. | 4.6e-09 | 0.08 | 1.4e-69 | 15.8 |

## 4.8. Appendix G: GWAS replication results

**Table 12:** Association between sequence variants and PAD for 4458 subjects from the replication set. **(A)** Genome wide significant sequence variants. **(B)** Sequence variants associated with structural MRI brain phenotypes that also associate with PAD. Abbreviations: confidence interval (CI), minor allele frequency (MAF).

| **(A)** |  |  |  |  |  |  |
| --- | --- | --- | --- | --- | --- | --- |
| rs Number | Position<br>(GRCh38) | Allele<br>(min/maj) | MAF (%) | Effect | 95% CI | *P* Value |
| rs2435204 | chr17:45910839 | G/A | 26.6 | 0.08 | (0.03, 0.13) | 2.1e-03 |
| rs1452628 | chr1:214966544 | T/A | 36.2 | -0.07 | (-0.11, -0.02) | 1.1e-03 |

| **(B)** |  |  |  |  |  |  |
| --- | --- | --- | --- | --- | --- | --- |
| rs Number | Position<br>(GRCh38) | Allele<br>(min/maj) | MAF (%) | Effect | 95% CI | *P* Value |
| rs2790099 | chr6:45475612 | C/T | 36.0 | -0.07 | (-0.11, -0.02) | 2.4e-03 |
| rs6437412 | chr3:194776533 | C/T | 28.2 | -0.05 | (-0.09, 0.00) | 4.6e-02 |
| rs2184968 | chr6:126439848 | C/T | 46.0 | 0.06 | (0.02, 0.10) | 2.9e-03 |

## 4.9. Appendix H: Effect of random CNN weight initialization on association between variants and association

**Table 13:** Correlation between UKB PAD (*N* = 12395) and the five reported sequence variants. Each subtable shows Pearson’s r correlation test results for five PADs (calculated by retraining our method five times) and a specific variant.

| **(A) Marker: rs2435204** |  |  |
| --- | --- | --- |
|  | Correlation (%) | P-Value |
| Original PAD | 6.0 | 2.1e-11 |
| PAD 1 | 5.0 | 2.1e-08 |
| PAD 2 | 4.9 | 4.6e-08 |
| PAD 3 | 5.2 | 6.6e-08 |
| PAD 4 | 5.4 | 1.5e-09 |

| **(B) Marker: rs1452628** |  |  |
| --- | --- | --- |
|  | Correlation (%) | P-Value |
| Original PAD | -5.5 | 7.1e-10 |
| PAD 1 | -4.6 | 3.3e-07 |
| PAD 2 | -4.3 | 1.9e-06 |
| PAD 3 | -4.3 | 1.7e-06 |
| PAD 4 | -3.8 | 1.6e-05 |

| **(C) Marker: rs2790099** |  |  |
| --- | --- | --- |
|  | Correlation (%) | P-Value |
| Original PAD | -4.2 | 3.8e-6 |
| PAD 1 | -2.5 | 5.6e-3 |
| PAD 2 | -2.8 | 2.0e-3 |
| PAD 3 | -3.0 | 9.7e-4 |
| PAD 4 | -2.4 | 7.1e-3 |

| **(D) Marke: rs6437412** |  |  |
| --- | --- | --- |
|  | Correlation (%) | P-Value |
| Original PAD | -3.8 | 2.6e-05 |
| PAD 1 | -3.5 | 9.0e-05 |
| PAD 2 | -2.7 | 2.4e-03 |
| PAD 3 | -3.5 | 9.8e-05 |
| PAD 4 | -3.2 | 3.7e-04 |

| **(E) Marker: rs2184968** |  |  |
| --- | --- | --- |
|  | Correlation (%) | P-Value |
| Original PAD | 3.7 | 3.6e-05 |
| PAD 1 | 2.6 | 3.6e-03 |
| PAD 2 | 2.9 | 1.0e-03 |
| PAD 3 | 2.2 | 1.2e-02 |
| PAD 4 | 2.6 | 3.4e-03 |

## 4.10. Appendix I: GTEx eQTL Analysis

**Figure 12:**
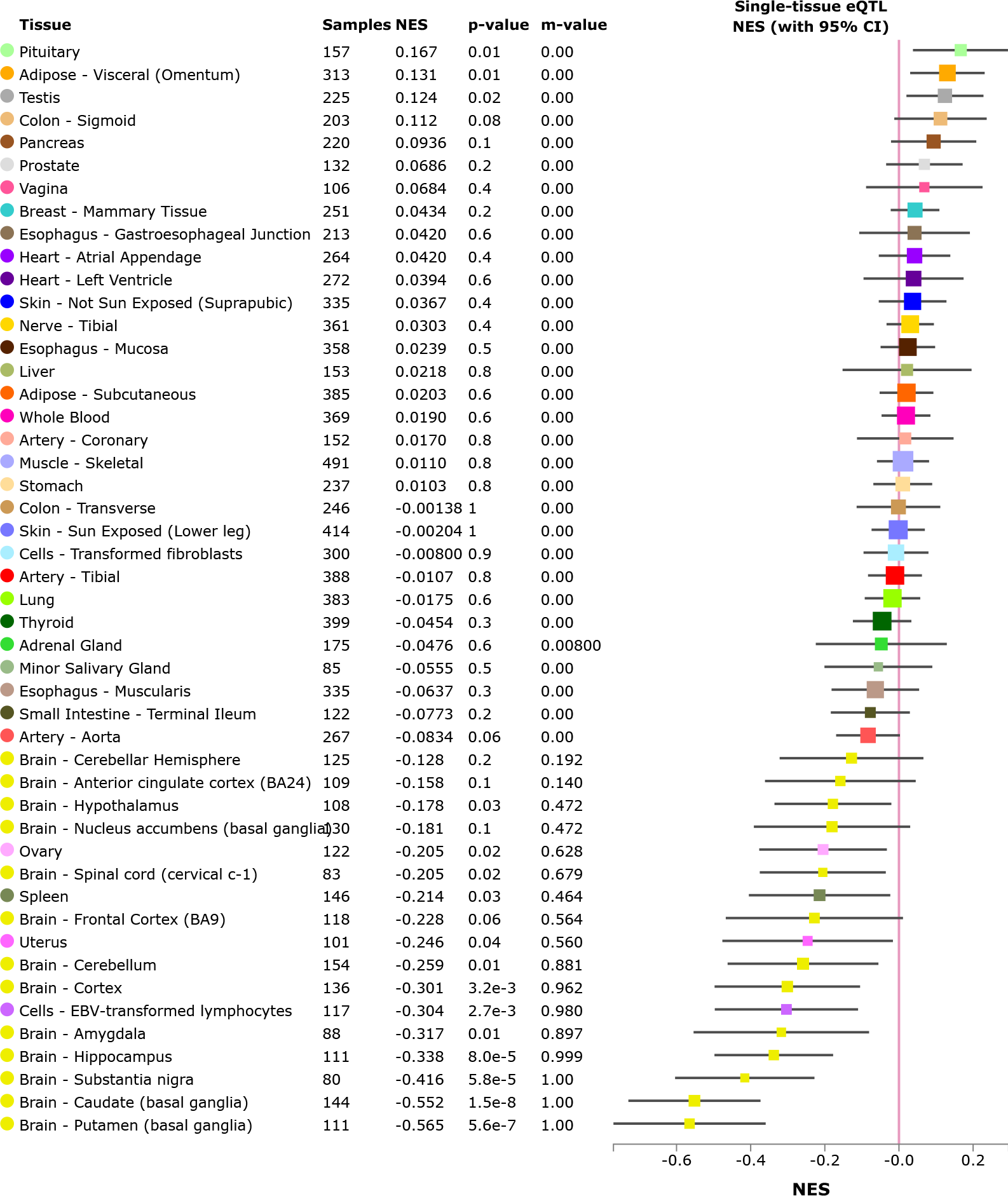
Multi-tissue eQTLs comparison for the *RUNX2* gene and rs2790099. Data Source: GTEx Analysis Release V7 (dbGaP Accession phs000424.v7.p2)

## Footnotes

1 Appendix D includes scatter plots of the CNN test set predictions against chronological age.

2 Appendix B includes more information and results about of the regression methods trained on extracted features.

3 http://brain-development.org/ixi-dataset/

4 In Appendix D the test set predictions before and after transfer learning are shown on a scatter plot against chronological age (Figure 11).

5 Appendix C gives a more detailed description of the tests.

6 Section 4.5 contains information about how the brain structure variants were identified.

7 Section 4.5 provides more information about the exact procedure.

8 Appendix F lists associations between the sequence variants and SBM/VBM brain structure pheno-types and correlation between PAD and the brain structure phenotypes (Tables 7–11).

9 PAD is adjusted for ICV, thus the observed effect on PAD is not caused by ICV.

10 www.ukbiobank.ac.uk/imaging-scanning-study/

11 Histograms of the age distribution of the three datasets mentioned are shown in Appendix A.

12 https://surfer.nmr.mgh.harvard.edu/fswiki/recon-all

13 www.neuro.uni-jena.de/cat/

14 https://cran.r-project.org/web/packages/ICC/index.html

15 GTEx Analysis Release V7 (dbGaP Accession phs000424.v7.p2)

16 http://biobank.ctsu.ox.ac.uk/crystal/field.cgi?id=20016)

17 http://biobank.ctsu.ox.ac.uk/crystal/field.cgi?id=20016)

18 http://biobank.ctsu.ox.ac.uk/crystal/label.cgi?id=100030

19 http://biobank.ctsu.ox.ac.uk/crystal/field.cgi?id=20018

20 http://biobank.ctsu.ox.ac.uk/crystal/field.cgi?id=20023

21 http://biobank.ctsu.ox.ac.uk/crystal/label.cgi?id=122

22 https://biobank.ctsu.ox.ac.uk/crystal/label.cgi?id=121

23 https://biobank.ctsu.ox.ac.uk/crystal/label.cgi?id=100077

